# Studying the Effect of Oral Transmission on Melodic Structure using Online Iterated Singing Experiments

**DOI:** 10.1101/2022.05.10.491366

**Authors:** Manuel Anglada-Tort, Peter M. C. Harrison, Nori Jacoby

**Affiliations:** Max Planck Institute for Empirical Aesthetics, Frankfurt am Main, Germany; University of Cambridge, Cambridge, UK

**Keywords:** iterated learning, singing, music, cultural evolution, transmission experiments

## Abstract

Since generations, singing and speech have been mainly transmitted orally. How does oral transmission shape the evolution of music? Here, we developed a method for conducting online transmission experiments, in which sung melodies are passed from one singer to the next. We show that cognitive and motor constraints play a profound role in the emergence of melodic structure. Specifically, initially random tones develop into more structured systems that increasingly reuse and combine fewer elements, making melodies easier to learn and transmit over time. We discuss how our findings are compatible with melodic universals found in most human cultures and culturally specific characteristics of participants’ previous musical exposure. Overall, our method efficiently automates online singing experiments while enabling large-scale data collection using standard computers available to everyone. We see great potential in further extending this work to increase the efficiency, scalability, and diversity of research on cultural evolution and cognitive science.

## Introduction

Singing – the vocal production of musical sounds – exists in every known human culture (Mehr et al., 2019; Nettl, 2010) and is likely to be amongst the first forms of communication and musical expression in human evolution (Tomlinson, 2015). Singing may have already been present in our closest ancient human relatives, Neanderthals (Mithen, 2006); and singing abilities emerge spontaneously in early child development (Dowling, 1999). At around 2 years of age, children start to generate recognisable songs and once they reach adulthood, most people can sing accurately in time and in tune (Dalla Bella, Giguère, & Peretz, 2007). Thus, singing is particularly suitable for studying the biological and cultural foundations of music evolution (Jacoby et al., 2019; Mehr & Krasnow, 2017).

Oral transmission is the main mechanism by which songs are passed through generations (Shanahan & Albrecht, 2019). In this simple act of transmission – hearing and singing back a song – it is likely that the singer introduces some variation to the new production, either accidentally or on purpose. Naively, one might expect that oral transmission just introduces random noise into the sung production. In practice, however, it is thought that oral transmission shapes musical systems in systematic ways that reflect human reproduction biases (Mehr, Krasnow, Bryant, & Hagen, 2020; Savage et al., 2022). With cultural exposure, people internalize these regularities and, in turn, are more likely to feature them in future productions. Thus, it is possible that the oral transmission of music in early humans resulted in the emergence of shared melodic features observed in musical traditions across the world (Savage, Brown, Sakai, & Currie, 2015; Mehr et al., 2019).

This paper focuses on transmission biases for melodies, including both cognitive constraints (categorical perception, memory limits, prior expectations; Burns, 1999; Greenspon & Pfordresher, 2019; Thompson, 2013) and motor constraints (certain music features are physically easier to produce than others; Miton, Wolf, Vesper, Knoblich, & Sperber, 2020; Tierney, Russo, & Patel, 2011). For example, memory biases facilitate the transmission of distinctive melodic features, such as the use of scales that are not uniformly symmetric (Pelofi & Farbood, 2021). Many regularities in melodies may also stem from the energetic costs associated with vocal production, such as the predominance of arch-shaped melodic contours or a bias towards small steps between adjacent notes (Huron, 2006; Savage, Tierney, & Patel, 2017).

Transmission chain experiments have proven particularly useful to study in the laboratory both perceptual priors and the evolution of cultural artifacts, such as language (Griffiths & Kalish, 2005; Langlois, Jacoby, Suchow, & Griffiths, 2021; Scott-Phillips & Kirby, 2010; Smith, Kirby, & Brighton, 2003; Thompson, Kirby, & Smith, 2016). Recently, researchers have begun to apply similar paradigms to the music domain, revealing the emergence of music regularities in rhythm (Jacoby & McDermott, 2017; Ravignani, Delgado, & Kirby, 2016) and melody (Lumaca & Baggio, 2017; Shanahan & Albrecht, 2019; Verhoef & Ravignani, 2021). However, performing such studies in complex production modalities such as singing is challenging. For example, covering the vast combinatorial musical space requires multiple transmission chains per experiment and the use of neutral sampling procedures that do not introduce bias on the music stimuli itself. Moreover, production tasks should be natural to participants (e.g., singing) and scalable to large-scale data collection.

Here we developed an automatic online pipeline to perform large-scale cultural transmission experiments in the singing modality. Participants are initially presented with a random “seed” melody (a sequence of pitches randomly generated from a continuous space) and asked to reproduce it by singing back (Figure 1A). Participants’ reproductions are then synthesized on the fly to create the stimuli for the next participants. Over the experiment’s generations, participants’ reproduction errors get amplified, reflecting transmission biases. Importantly, our method does not assume culturally specific knowledge about scale systems *a priori* – e.g., using discrete musical systems such as the Western 12-tone equal temperament scale. Instead, we randomly sample melodies from a continuous pitch space. Consequently, our method is hypothesis-neutral and applicable to individuals from any musical or cultural background.

**Figure 1:**
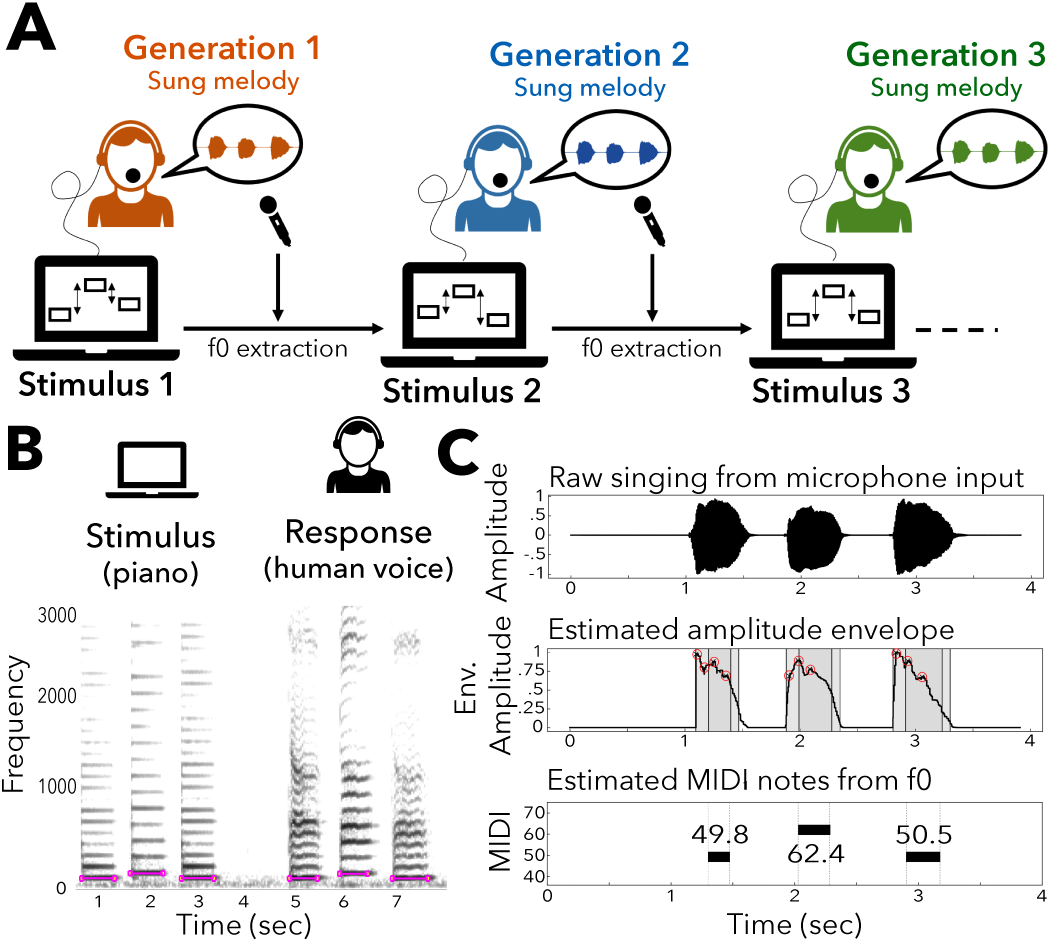
Online Iterated Singing Paradigm. (A) Participants hear sequences of tones generated by a computer and reproduce them back by singing. Vocal reproductions are synthesized online and played to the next participant as the input melody. (B) Spectrogram showing a three-tone melody and corresponding vocal reproduction. (C) Singing extraction: we input the recording from the microphone and estimate MIDI notes using f0 extraction techniques.

We validate this method in two online experiments with short (three-tone) and longer (five-tone) melodies. We show that transmission biases in singing can give rise to melodic structural features that are consistent with statistical universals – i.e., features that occur frequently in musical traditions across the world (Savage et al., 2015). In particular, we find that (1) melodies are biased towards a small vocabulary of intervals (pairs of notes), (2) increasingly exhibit arch-shaped musical contours (the pitch sequences of ups and downs), and (3) are composed of small intervals (less than a perfect fifth). Parallel to this, evolving melodies tend to align with melodic pleasantness (estimated by a separate rating experiment with Western listeners) and musical exposure (estimated by a large corpus of popular Western melodies). As a result, melodies become increasingly easier to learn and transmit. We conclude by discussing the implications and limitations of these findings.

## Methods

### Participants

We recruited a total of 186 participants that provided consent in accordance with the Max Planck Society Ethics Council approved protocol (application 2020-05). All participants were recruited online using Amazon Mechanical Turk (AMT) with the following three constraints on recruitment: (i) participants must be from the US, (b) at least 18 years old, and (iii) have a 95% or higher approval rate on previous tasks on AMT. Participants were paid at a US $9/hour rate according to how much of the experiment they completed. The completed experiments took approximately 20-25 minutes.

### Automated Online Implementation

The experiments were implemented in PsyNet (https://www.psynet.dev/), a Python package for performing complex online behavioral experiments at large scale (e.g., Harrison et al., 2020). Participants interact with the experiment via a Chrome web browser, which communicates with a backend Python server cluster responsible for organizing the experiment and collecting data.

### Singing Extraction

We use a three-step automated process to estimate the fundamental frequency (f0) of notes in vocal productions (Figure 1B). First, we clean the audio signal by applying a bandpass filter with cutoff frequencies of 80-6000 Hz. Second, we apply an autocorrelation-based pitch estimation algorithm to extract f0 pitch from sung segments (Boersma, 1993), implemented using parselmouth (Jadoul, Thompson, & de Boer, 2018), a Python interface to access Praat. Finally, we identify voiced segments (continuous time spans with reliably f0 tracking), and compute the median f0 for each segment (Figure 1C).

### Melody Generation

We parameterize the space of melodies as lists of numbers in a logarithmic scale, using MIDI (Rothstein, 1992), where a MIDI note (*m*) represent a frequency of 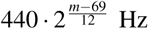 Hz. To generate an initial random melody, we first assign a comfortable singing register (low vs high) to each participant based on their self-reported gender or vocal range. Next, we obtain a reference pitch for each melody by uniformly sampling a real number within a roving width of ±2.5 semitones around the center of the singing register (52.5 for low-register and 63.5 for high-register). Roving the reference pitch helps minimize carryover effects between trials, where a given trial is interpreted in terms of an implied tonal context from the previous trial. Finally, we sample each pitch in the melody from a uniform distribution with a width of ±7.5 semitones centered on the reference pitch.

All tones were 300 ms long in duration and presented with an inter-tone interval of 800 ms. Tones were played using a piano timbre via Tone.js, a Web Audio framework for generating sound in the browser (https://tonejs.github.io/).

### Procedure

We used a combination of techniques to ensure high data quality when recruiting participants online in complex production experiments (Anglada-Tort, Harrison, & Jacoby, 2022), including audio calibration and recording tests, a prescreening task to ensure headphones use (Woods, Siegel, Traer, & McDermott, 2017), and a singing performance test to measure participants singing accuracy and exclude both fraudulent participants and computer bots.

In the main singing task, participants completed a total of 50 trials, which consisted of hearing a melody and imitating it by singing back each note to the syllable ‘Ta’. In each trial, participants were randomly allocated to one of the parallel transmission chains. Each participant could only contribute once to each chain, thereby contributing to a maximum of 50 different chains per experiment. After each trial, we analyzed participants’ recordings (see Singing Extraction) to determine whether we could detect the requested number of notes per trial. Trials that did not satisfy this condition failed and did not contribute to the chain. In such cases, new participants were allocated to that trial until a valid response was given.

## Results

### Three-Note Melodies

Experiment 1 examined short melodies composed of two intervals (3 notes, Figure 2A). This stimulus space can be defined along two continuous dimensions (i.e., interval 1 and interval 2), where integer locations correspond to the Western 12-tone equal temperament scale. We explored this space with 333 across-participant chains with 10 generations per chain (3,330 singing trials). A total of 57 US participants contributed to this experiment (aged 18-63, *M* = 38.95, *SD* = 13.31; 41.1% with low and 58.9% with high vocal range).

**Figure 2:**
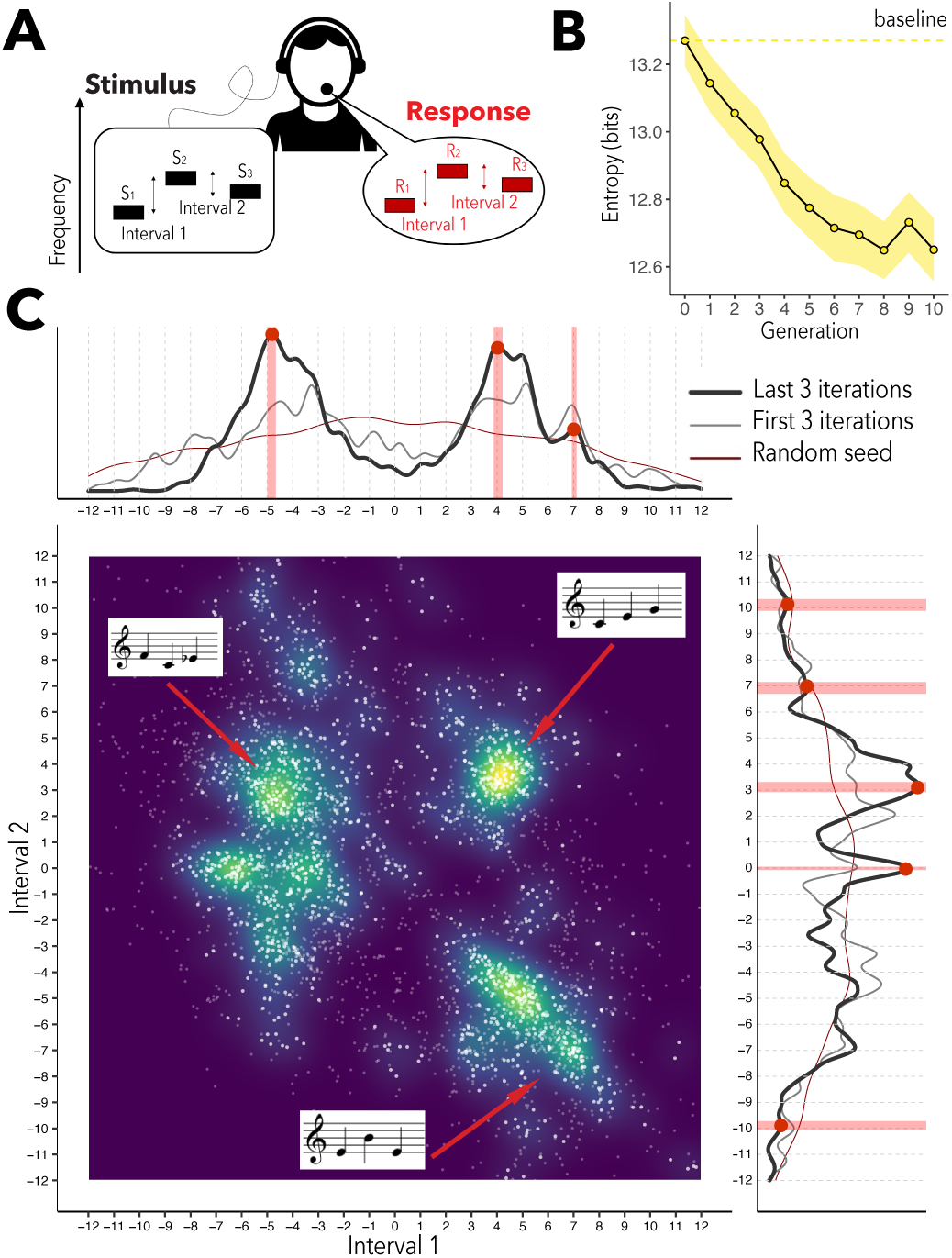
Iterated Singing Experiment with Short Melodies. (A) Participants hear and sing back three-note melodies. (B) The entropy of the distribution of intervals over generations, suggesting an increase in combinatorial structure. (C) Two dimensional KDE over the last three generations. On the top and right panels we plot the distribution of each interval separately (dark grey line), including the distribution of the random initial set of melodies (red line) and the first three generations (light grey line). Statistically significant peaks (90% CI) are indicated by the red dots and shaded areas. *Note*. Error bars represent bootstrapped standard error (1000 replicates).

Figure 2C shows the aggregated results of the iterated singing experiment using a kernel density estimator (KDE) over the locations of the reproductions^1^. Despite making no assumptions about musical systems *a prori*, we see that productions in the last generations are concentrated in few locations, displaying a rich structure that is consistent with Western discrete scale systems. For example, the KDE reveals peaks at common and prototypical interval combinations in Western music (top-right quadrant in Figure 2C): the major scale, featuring melodies with consecutive intervals at [4, 3], [5, 4], and [5, 3]. Another popular area (bottom-right quadrant) consists of arch-shaped musical contours (melodies that first ascend in pitch and then descent), mostly peaking in the perfect fourth [5, -5] and perfect fifth [7, -7]. We also computed the marginal distribution of the first and second interval separately (see top and right panels in Figure 2C). Interestingly, the two intervals have clearly distinct distributions, suggesting that they play different roles in two-note melodies.

Next, we used MATLAB’s *findpeaks* method to identify significant peaks that correspond to integer semitones categories. To measure confidence, we repeated this analysis in 1,000 bootstrap datasets (sampling with replacement over chains), identifying each time those peaks that fell into integer semitone categories (±0.5 around each integer). We then calculated the probability of having at least one peak associated with each integer category over all bootstrap datasets. Statistical significant peaks close to integer semitones are indicated in Figure 2C by the red dots and shaded areas, representing 90% confidence intervals (CI). The observed peaks are consistent with the notion of consonance in Western music (Halpern & Bartlett, 2010) – e.g., peaks in melodic intervals that maximize consonance, such as the major third (4), perfect fourth (5), and perfect fifth (7).

To assess whether the structure of the distribution of intervals increased over generations, we used Shannon’s entropy (Shannon, 1948), *H* = ∫*p*(*x, y*) log *p*(*x, y*) *dx dy*, where *p*(*x, y*) is estimated from the 2D KDE of Figure 2C. Entropy measures the extent to which the distribution is random or concentrated in particular places in the space (forming “peaks” or “structure”). As shown in Figure 2B, entropy decreases significantly over generations (*B* = -.061, 95% CI [-.077, - .045])^2^. This suggests that melodies in later generations are composed of a smaller vocabulary of intervals that are increasingly reused.

### Five-Note Melodies

Experiment 2 examined longer melodies composed of four intervals (five notes, Figure 3A). The resulting stimulus space comprises four dimensions (one for each interval). We explored this space with 120 across-participant chains with 10 generations per chain (1,200 singing trials). A total of 38 US participants contributed to this experiment (aged 20-65, *M* = 38.94, *SD* = 12.48; 50.51 % with low and 49.49% with high vocal range). As with three-note melodies, there was a decrease in entropy^3^ over generations (*B* = -.235, 95% CI [-.273, -.197]), suggesting an increase in combinatorial structure (Figure 3B).

**Figure 3:**
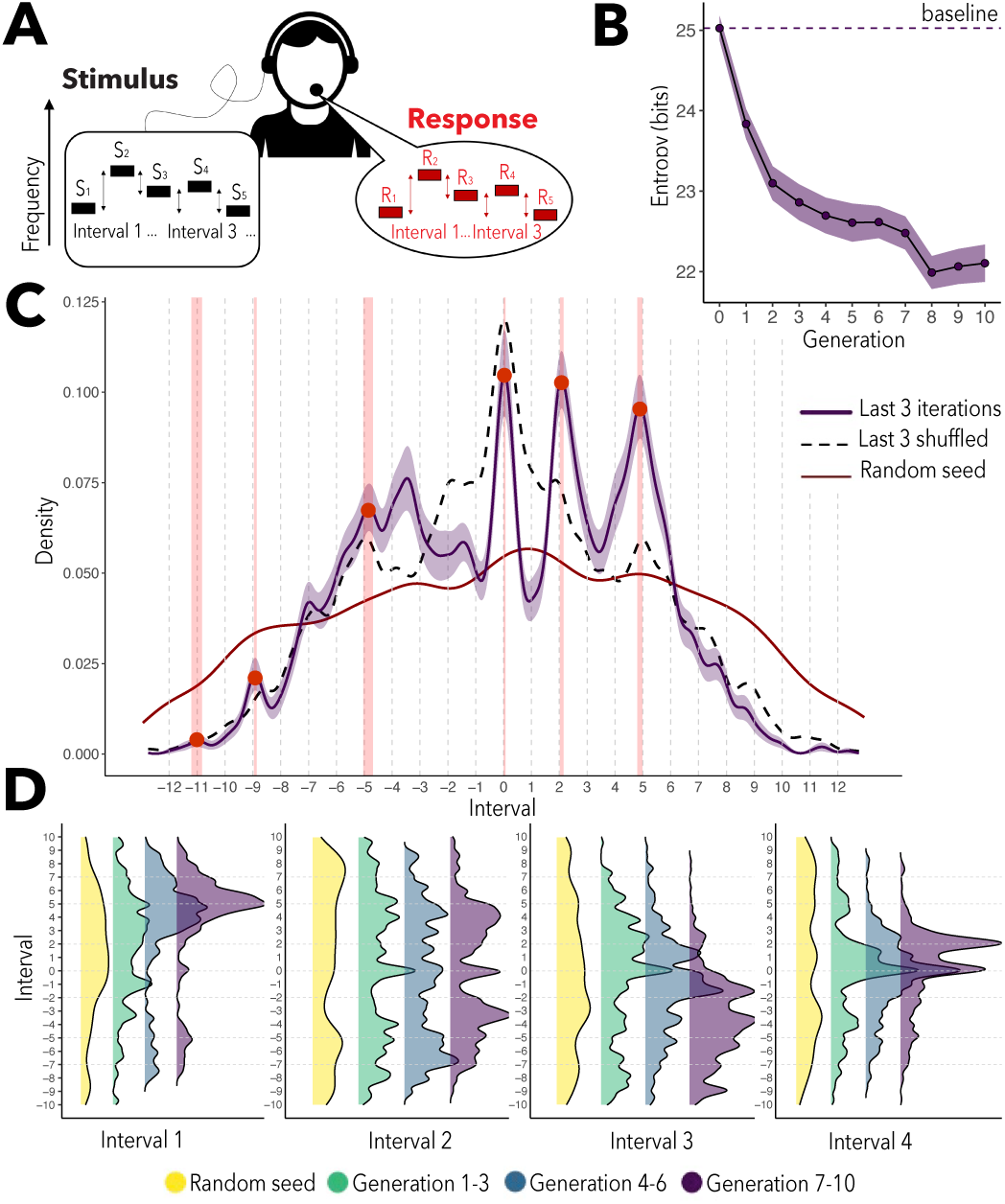
Iterated Singing Experiment with Longer Melodies. (A) Participants hear and sing back five-note melodies. (B) The entropy of the distribution of intervals over generations, suggesting an increase in combinatorial structure. (C) KDE of the four melodic intervals in the last three generations (purple line). We also plot the distribution of the initial random stimulus set (red line) and the sung reproductions in the last three generations shuffling the order of the pitches in each melody (black dashed line). (D) KDEs of the four intervals plotted separately over generations. *Note*. Error bars represent bootstrapped standard error (1000 replicates).

Figure 3C shows the results of the iterated singing experiment by plotting the distribution of the four intervals together in a 1-dimensional KDE. We identified 6 statistically significant peaks close to integer semitones, indicated in Figure 3C by the red dots and shaded areas representing 90% CI. In addition, we plot the distributions of the initial stimulus set (red line) and the reproductions in the last 3 generations but randomizing the order of the pitches in each melody 10 times (black dashed line). Comparing these distributions emphasizes the effect of sequential memory effects and preferences for certain melodic intervals, such as avoidance to the semitone (1) and attraction to the perfect fourth (5). Furthermore, Figure 3D shows the evolution of each interval separately, indicating that melodic intervals play different roles depending on their position in the melody. For example, five-tone melodies tend to start with a relatively large ascending interval at around 5 semitones (perfect fourth), and end with a small ascending interval at around 2 semitones (major second) or a unison.

### Effects of Oral Transmission on Melodies

Having demonstrated that our method is efficient to explore large melodic spaces using online iterated singing experiments, we next look at the effects of oral transmission on structural features of melodies. This search was motivated by two intertwined bodies of research: (1) cross-cultural studies looking at widespread melodic features across cultures (Brown & Jordania, 2013; Mehr et al., 2019; Savage et al., 2015, 2017) and (2) research on culturally specific characteristics of listeners’ previous musical exposure (Hannon & Trehub, 2005; Loui, Wessel, & Kam, 2010; Jacoby et al., 2019, 2021). While the former suggests that knowledge of musical structure may stem from universal constraints in our vocal and auditory system, the latter suggests that such knowledge may be acquired via exposure to specific music cultures. Although the current experiments are unable to distinguish between these two accounts, one should consider both when interpreting the following results.

Figure 4 summarizes the results of this analysis. First, we found that melodies are biased towards a small homogenized vocabulary of intervals (Figure 4A). This analysis was performed using the peak finding procedure described above. Indeed, the number of detected peaks significantly decreases over generations (Experiment 1: *B* = -.282, 95% CI [-.448, -.115]; Experiment 2: *B* = -.311, 95% CI [-.481, -.140]). This finding is consistent with large-scale quantitative data showing that melodies across cultures tend to contain a small number of elements (7 or less) per octave (Savage et al., 2015).

**Figure 4:**
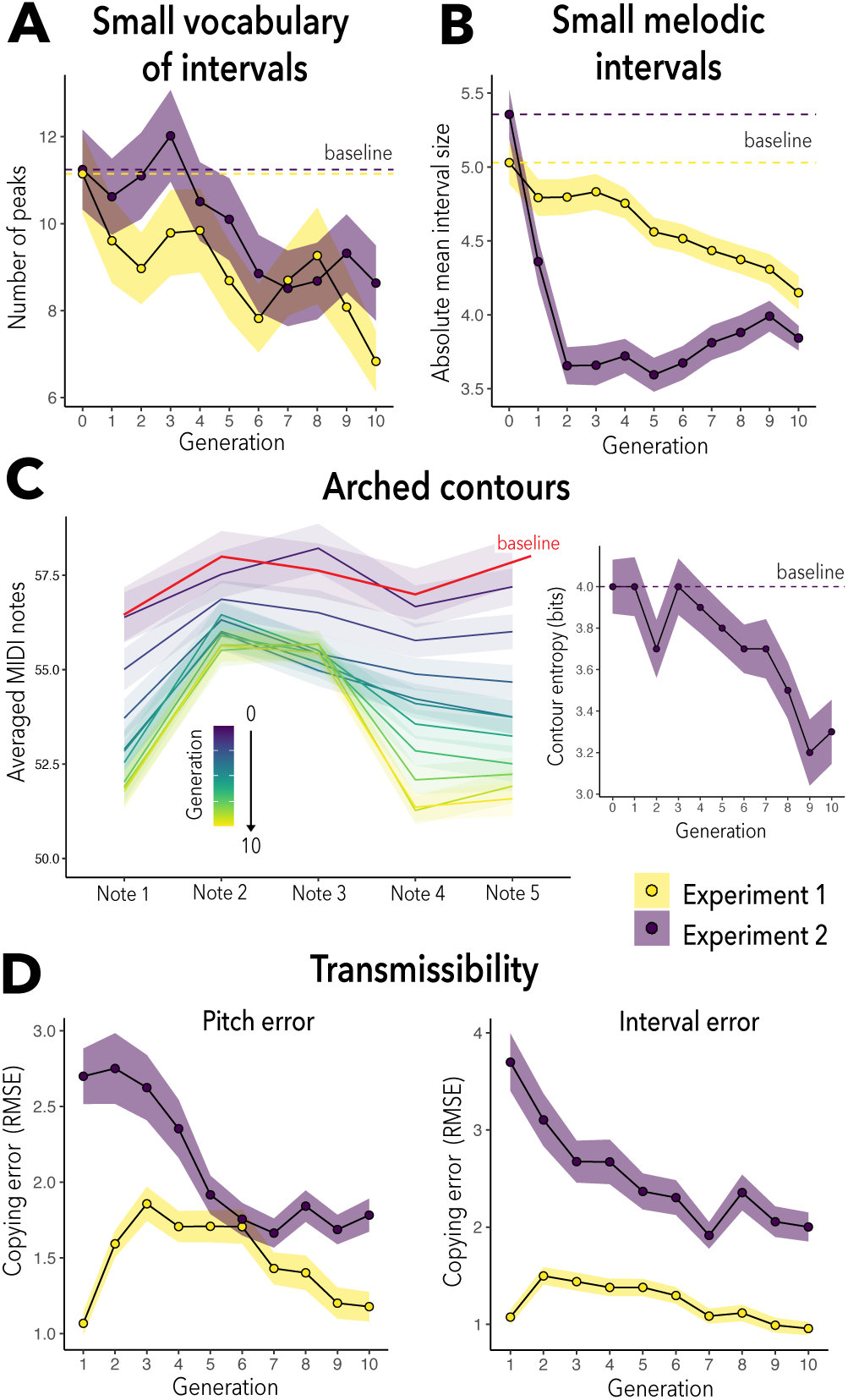
Effects of Oral Transmission on Melodies. (A) Melodies are biased towards a homogenized small vocabulary of intervals. (B) Melodic intervals become smaller over generations. (C) Melodies tend to use arched contours (left plot); fewer musical contours are increasingly reused and combined over generations (right plot). (D) Melodies become easier to learn and transmit. *Note*. All error bars represent bootstrapped standard error (1000 replicates).

Second, we observed that melodic intervals become significantly smaller over generations (Figure 4B), as indicated by the absolute mean interval size (Experiment 1: *B* = -.078, 95% CI [-.112, -.045]; Experiment 2: *B* = -.074, 95% CI [-.111, -.037]). This effect is consistent with a widely observed feature that characterizes animal and human vocalizations (Savage et al., 2015; Tierney et al., 2011): jumps between consecutive pitches in vocal utterances tend to be small.

Third, orally transmitted melodies increasingly exhibit arch-shaped musical contours. Figure 4C (left plot) shows the average pitch of each note in the five-note melodies experiment, showing that melodies evolve from flat melodic contours (see baseline) to distinctive arched contours^4^. We further examined the emergence of archetypal contours by calculating a measure of contour smoothness using Shannon entropy. Each interval was coded to three possible values: 0 if it was approximately a unison (between -0.5 and 0.5 semitones), +1 if it was ascending (larger than 0.5 semitones), and -1 if it was descending (smaller than -0.5 semitones). We then computed the entropy of this discrete distribution of contour types over generations. The results indicate a significant decrease of contour entropy over generations (right plot in Figure 4C), revealing that fewer melodic contours become increasingly reused over time (*B* = -.074, 95% CI [-.100, -.047]). The prevalence of simple and archetypal contours has been found to be widespread both in bird song (Tierney et al., 2011) and human song across cultures (Savage et al., 2017).

Since our participants had significant prior exposure to Western music (they were all recruited from the US), we asked whether such exposure may have had an effect on evolving melodies. First, we looked at the interval distribution of a large corpus of popular Western melodies as a proxy of participants’ previous musical exposure (magenta color in Figure 5A). This corpus consisted of a a subset of 6200 melodies from the Lakh MIDI Dataset (Raffel, 2016) that have been matched to entries in the Million Song Dataset (Bertin-Mahieux, Ellis, Whitman, & Lamere, 2011)^5^. We then calculated how aligned the melodic intervals in our experiments were with the prevalence of intervals in the corpus. Results indicate that melodies become increasingly more aligned with the corpus data (Figure 5B, left plot), although this trend was only significant in Experiment 1 (*B* = .035, 95% CI [.010, -.059]; Experiment 2: *B* = .018, 95% CI [-.002, .04]).

**Figure 5:**
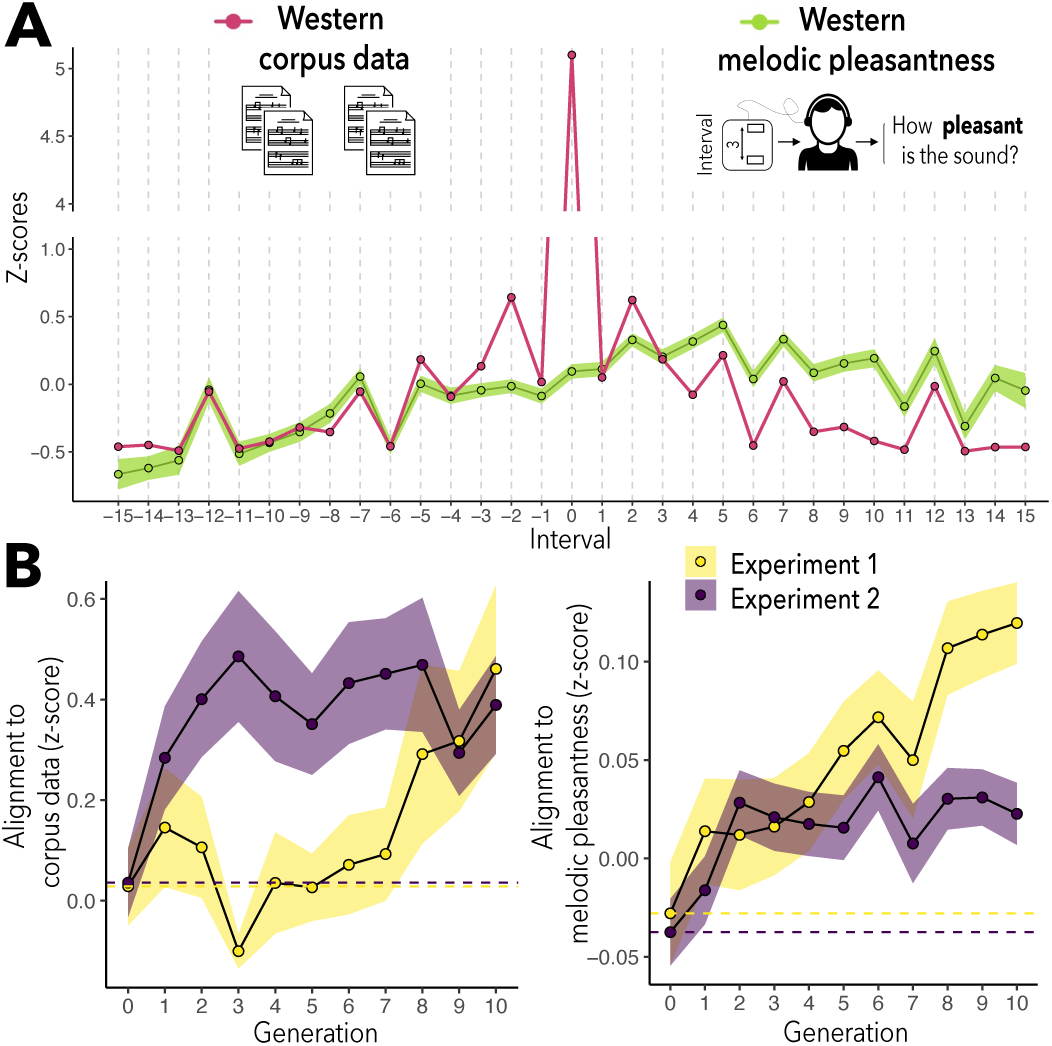
Alignment with Western Tonal Exposure. (A) Interval distribution of a large corpus of popular Western melodies (magenta) and perceived melodic pleasantness measured by a separate group of Western listeners (green). (B) Melodies become increasingly more aligned with Western corpus data (left plot) and perceived melodic pleasantness (right plot). *Note*. All error bars represent bootstrapped standard error (1000 replicates).

In addition, we ran a listening experiment with a separate group of Western participants to measure the perceived pleasantness of melodic intervals. In this experiment, 91 US participants (aged 22-77, *M* = 38.83, *SD* = 11.90) were presented with melodic intervals (two tones played sequentially) and asked to rate how pleasant they sounded, on a 7-point scale. We evaluated 5000 intervals ranging from -15 to 15 semitones, with each participant rating an average of 80 intervals. The intervals were generated following the same procedure described above (see Melody Generation). However, to reduce the vast pitch space, this experiment only used integer MIDI notes. The resulting aggregated scores provide a measure of melodic pleasantness in Western music (green color in Figure 5A). As shown in Figure 5B (right plot), there is a small but significant increase in the perceived pleasantness of melodic intervals over generations (Experiment 1: *B* = .014, 95% CI [.008, .019]; Experiment 2: *B* = .004, 95% CI [.001, .007]).

Finally, we examined the evolution of copying error (the distance between the target and sung production), using the root mean square error (RMSE) in both pitch and interval domains (Figure 4D). In Experiment 1, we excluded the first generation from the analysis because copying error was surprisingly small (this is probably an artifact of the melody generation process). However, after the first generation, copying error significantly decreased over time (Pitch: *B* = -.075, 95% CI [-.101, -.049]; Interval: *B* = -.072, 95% CI [-.094, -.049]). In Experiment 2, the decrease in copying error was clear from the start and comparatively larger than in Experiment 1 (Pitch: *B* = -.132, 95% CI [-.176, -.088]; Interval: *B* = -.161, 95% CI [-.221, -.100]). Overall, this finding suggests that oral transmission shapes structural features of melodies so they become easier to learn and reproduce over generations.

## Discussion

We introduced a novel method to automate cultural transmission experiments in the singing modality and over the internet. Overall, we found that oral transmission has profound effects on the evolution of melodies, shaping initially random sounds into more structured systems – i.e., using fewer building blocks (intervals, contours) that are increasingly reused and combined. Importantly, structural features that emerged artificially from our experiments are largely consistent with widespread melodic features observed in most musical traditions across the world (Brown & Jordania, 2013; Mehr et al., 2019; Savage et al., 2015). Moreover, we found that melodies tend to align with certain cultural conventions of Western music (e.g., use of harmonic intervals), reflecting our participants’ predominantly Western background.

How can we explain the emergence of such melodic features? They likely stem from a complex interplay between motor constraints, cognitive biases, and cultural exposure (Tomlinson, 2015). However, the current paradigm is limited in its ability to disentangle different kinds of transmission biases, such as cognitive versus production constraints. Despite this, we found that melodies in our experiments were increasingly more aligned with a separate perception-only preference experiment (Figure 5B), suggesting that perception (and not only production) influences the oral transmission of music. Future experiments will help us better determine the relative contribution of perception and production, for instance by conducting perceptual only experiments on cultural transmission (Harrison et al., 2020).

Furthermore, our participants had significant prior exposure to Western music and, therefore, the current design is unable to distinguish universal transmission biases (e.g., stemming from biological factors) from cultural transmission biases (e.g. stemming from familiarity with Western tonality). Future experiments can systematically test this by conducting large-scale cross-cultural experiments with participants from diverse musical backgrounds (Jacoby et al., 2021).

Above all, our method is useful for recovering many transmission biases within a single coherent paradigm that is cross-culturally generalizable. In particular, our method makes no assumptions about musical systems *a prori*, is natural and intuitive to everyone (e.g., singing), and works using standard computers available to most people online, enabling experiments that would be nearly impossible in the laboratory. We are truly excited about the potential of this work to increase the reach, scalability, and diversity of research on cultural evolution, music cognition, and cognitive science.

## Acknowledgments

We thank Luke Poeppel for helping develop the singing extraction technology and the members of the Computational Auditory Perception Group for their support and feedback. We also thank the artists *PIXARTIST* for the design of the listening icon (https://www.flaticon.com/premium-icon/listening_5725503) and *Freepik* for the design of the music score icon (https://www.flaticon.com/free-icon/score_92971).

## Footnotes

1 The bandwidth in the 2-dimensional KDE was estimated using Scott’s method (Scott, 2015). In all other analyses, including 1-dimensional KDEs, the bandwidth was set to 0.25 (a quarter tone).

2 All trend analyses in this paper are analyzed using linear regressions, with 95% confidence intervals derived by bootstrapping over chains (1,000 replicates, Gaussian approximation).

3 Entropy for this experiment was approximated by summing the contributions from the entropies of all consecutive intervals, where the entropy for each consecutive interval was computed from the 2-dimensional KDE (as we did for the three-tone melodies).

4 This analysis only uses data from Experiment 2 because the proportion of musical contours in Experiment 1 is skewed from the start due to the sampling procedure.

5 This dataset provides a collection of audio features and metadata for a million contemporary popular music tracks (http://millionsongdataset.com/).

